# A mutant of the phototoxic protein KillerRed that does not form DsRed-like chromophore

**DOI:** 10.1101/868877

**Authors:** Dmitry A. Gorbachev, Karen S. Sarkisyan

## Abstract

We report a mutant of the phototoxic fluorescent protein KillerRed protein with fluorescence in the green part of the spectrum. The mutant variant carries mutations I64L, D114G, and T115S and does not form a DsRed-like chromophore. The protein can be used as a template to create new genetically encodable photosensitizers that are spectrally different from KillerRed.

## Introduction

Fluorescent proteins are widely used as genetically encodable tags for optical labeling of living systems [1]. Their chromophores are located inside the protein structure and are protected from the surrounding solvent; therefore, most of the existing fluorescent proteins are passive reporter molecules: irradiation with light does not significantly affect cells expressing these markers.

At the same time, a unique family of genetically encoded photosensitizers has been developed based on the fluorescent protein anm2CP. Upon light illumination, members of the family produce reactive oxygen species that can damage the cell [2]. Structural studies of these proteins have identified a water-filled channel that connects the chromophore to the solvent. This structural feature of phototoxic fluorescent proteins is thought to be responsible for the efficient diffusion of reactive oxygen species into the environment [3,4].

KillerRed was the first protein engineered to produce reactive oxygen species demonstrating phototoxicity levels exceeding other fluorescent proteins more than thousand-fold [2]. Depending on the cellular localization and the excitation light dose, reactive oxygen species generated by KillerRed can lead to various physiological consequences — from inactivation of fusion proteins [2], to cell division arrest [5,6] or cellular death through necrosis or apoptosis [2,7]. Due to these capabilities, KillerRed is used as an optogenetic tool in cell biology to inactivate proteins with light, to study intracellular oxidative stress or to ablate specific cell populations. KillerRed has also been used as a photosensitizer for treatment of tumors in model systems [7–9].

Other genetically encoded photosensitizers have been created based on KillerRed, including SuperNova, a monomeric KillerRed variant with similar spectral characteristics [10], as well as orange fluorescent protein KillerOrange [11] and green fluorescent protein SuperNova Green [12]. Besides that, non-fluorescent protein-based photosensitizers miniSOG, Pp2FbFP and others that generate singlet oxygen have been developed [13,14].

None of the existing photosensitizers with green fluorescence can be universally used in relevant applications. Some proteins demonstrate incomplete or slow chromophore maturation rate; others may not generate the type of reactive oxygen species suitable for a particular application, or their phototoxicity may depend on the availability of external chromophore [15]. In this work, we aimed to find a mutant version of KillerRed that did not form a DsRed-like “red” chromophore and, therefore, could serve as a basis for the development of the new generation of efficient photosensitizers with green fluorescence.

## Material and methods

DNA amplification was performed using the Encyclo PCR Kit (Evrogen) on a PTC-200 Thermal Cycler (MJ Research). The analysis of amplification products was carried out in 1-2% agarose gel. Ethidium bromide was used at a concentration of 0.5 μg/ml.

Mutagenesis was performed by error-prone PCR. We added manganese ions and a skewed ratio of nucleotide triphosphates (50x mix: 0.2 MM dGTP, 0.2 MM dATP, 1 MM dCTP, 1 MM dTTP) into the PCR reaction mixture leading errors in Taq-polymerase-based DNA amplification. The average mutation rate was 8 nucleotide substitutions per 1000 amplified nucleotides after 25 cycles of PCR.

For electroporation, the ligation mixture was purified on Cleanup Mini DNA purification columns (Evrogen). 40 μl of electrocompetent cells were thawed on ice, and up to 5 μl of purified ligation mixture was added to thawed cells. Cells were then transferred into a pre-cooled electroporation cuvette (Bio-Rad, USA) and electroporated on the MicroPulser device (Bio-Rad, USA). Immediately after electroporation, 3 ml of SOB medium was added to the cuvette, and the bacterial suspension was transferred into 1.5 ml plastic tubes. The tubes were incubated for one hour in an incubator at 37°C and then plated on agar. The plates were incubated at 37°C for 18 hours. The average density of E. coli colonies was 5,000 per plate and the total diversity of genotypes in the library was estimated to be around 100,000 clones. The number of fluorescent colonies was 22%.

### Expression and purification of recombinant proteins

E. coli XL1 Blue was grown in 800 ml flasks in LB medium with ampicillin (100 mg/ml), induced with isopropyl-β-D-1-thiogalactopyranoside to the final concentration 0.5 mM and incubated for 3 hours. All further operations were performed on ice. The culture was centrifuged, the supernatant was discarded, the pellet was resuspended in 4 ml of phosphate buffer (pH 7.4), the suspension was lysed in Sonics Vibra Cell sonicator and centrifuged again. The supernatant was transferred into a new tube with 400 μl of Talon metal-affinity resin (Clontech), equilibrated with phosphate buffer. The tube was placed in a shaker for one hour at 200 rpm at room temperature. Then, the resin with the protein was washed several times with phosphate buffer and eluted with phosphate buffer containing imidazole (250 mM).

## Results

We relied on random mutagenesis to find KillerRed mutant with green fluorescence. The mutant library was cloned into pQE-30 vector, transformed into E. coli cells and grown on agar plates without induction. We visually screened bacterial colonies exposed by 400 nm and 480 nm light to identify mutants with significant green fluorescence and identified the KillerRed I64L/D114G/T115S mutant (**Table 1**, the numbering of positions in the protein is indicated according to the established notation so that the chromophore-forming residues are in positions 65-67 [1]). This mutant had very dim fluorescence in the red part of the spectrum while being noticeably fluorescent in green under when illuminated with 480nm light. Observed spectral properties indicated that the protein formed the “classical” GFP-like chromophore instead of the DsRed-like chromophore found in the parental KillerRed.

**Table 1.**
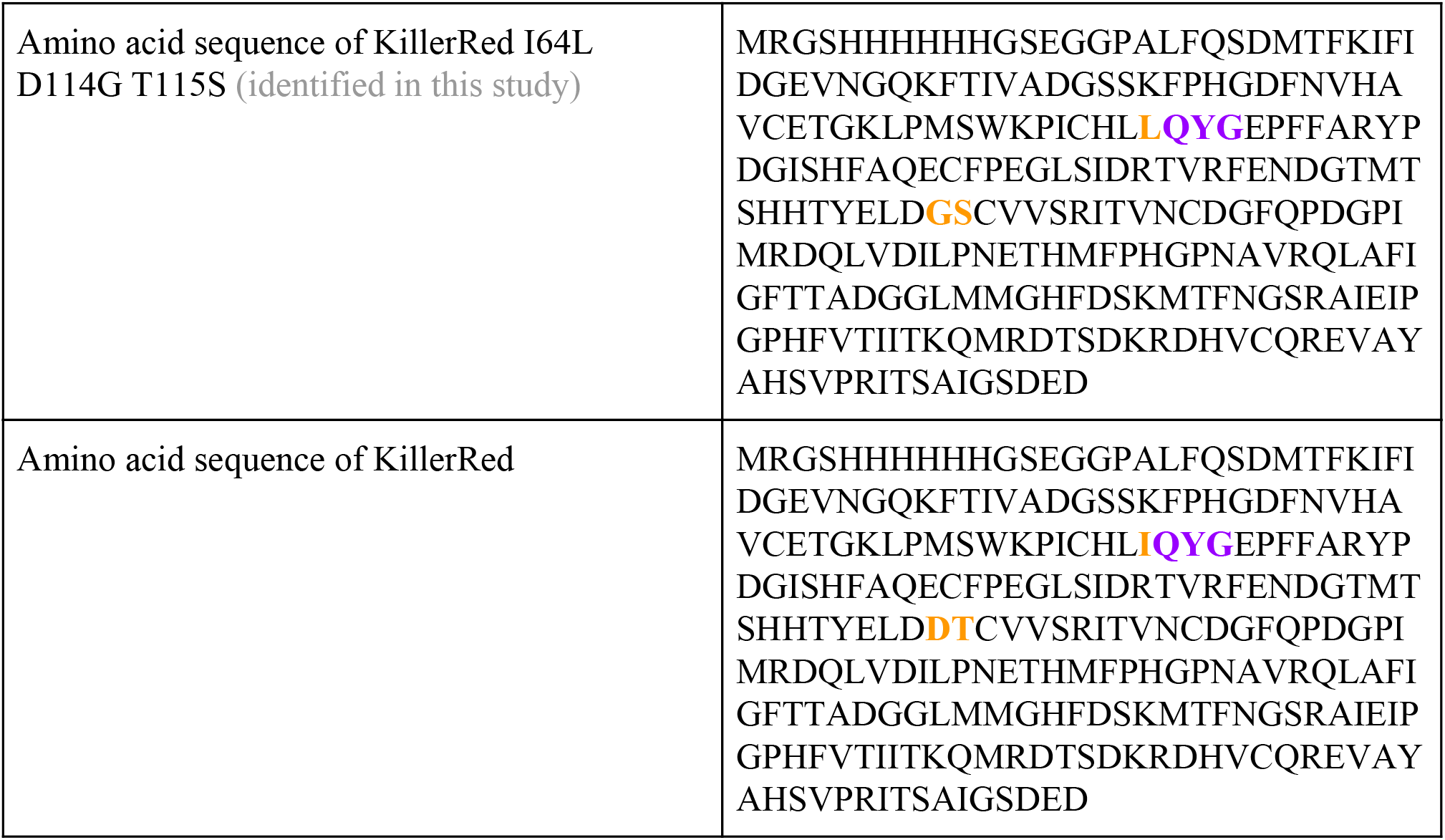
Amino acid sequences of proteins described in this study. Chromophore-forming residues are highlighted in purple, positions containing mutations are highlighted in orange.

Since proteins with GFP-like chromophores and DsRed-like chromophores have specific absorption spectra that are easily distinguishable from each other, we isolated and purified KillerRed protein and it’s mutant KillerRed I64L/D114G/T115S (Figure 1). The absorption spectrum of purified KillerRed I64L D114G T115S was significantly different from the absorption spectrum of KillerRed and had a pick with the maximum at 514 nm and a blue-shifted shoulder characteristic of GFP-like chromophores.

**Figure 1.**
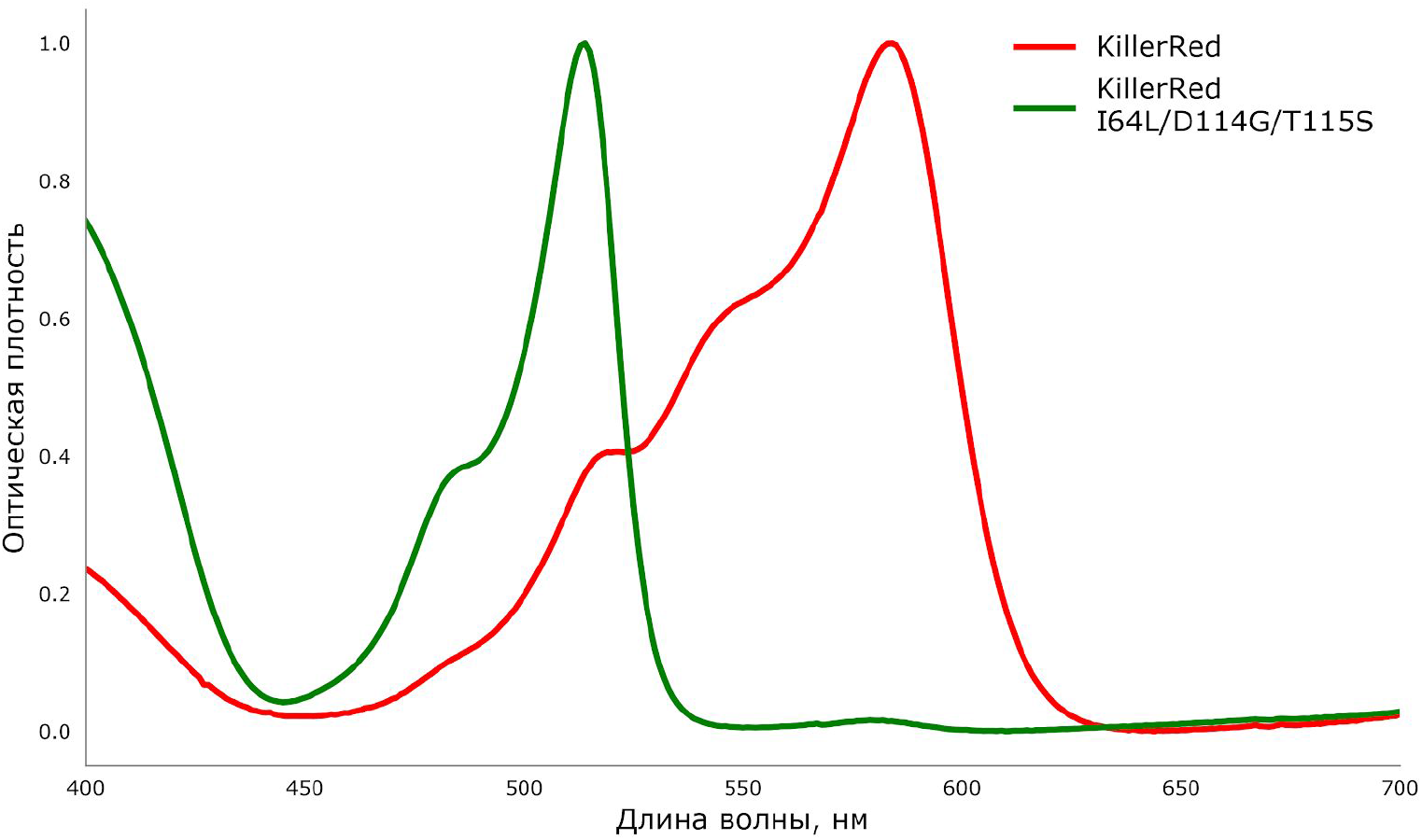
Absorption spectra of purified proteins KillerRed and KillerRed I64L/D114G/T115S.

## Discussion

The lack of absorbance at 550-600 nm in KillerRed I64L/D114G/T115S indicated that introduced mutations almost entirely prevented formation of DsRed-like chromophore. The peak with a maximum at 514 nm and a characteristic shoulder at 480–485 nm suggested that chromophore catalysis stopped at a “classical” GFP-like chromophore [1].

Mutations found in the KillerRed variant we identified are interesting in the context of existing literature on fluorescent proteins mutagenesis. In particular, mutations at position 64 were previously described as affecting chromophore maturation: for instance, in *Aequorea victoria* GFP, the F64L mutation improves maturation of the chromophore when expressed at 37C [1], while in the chromoprotein from *Acropora millepora* the S64C mutation changes the color of the protein [16]. Mutations D114G and T115S are located in the neighboring positions on the loop connecting the beta-strands 5 and 6 (**Figure 2**) and may contribute to the adaptation of the beta-barrel to the replacement of isoleucine with leucine at position 64.

**Figure 2.**
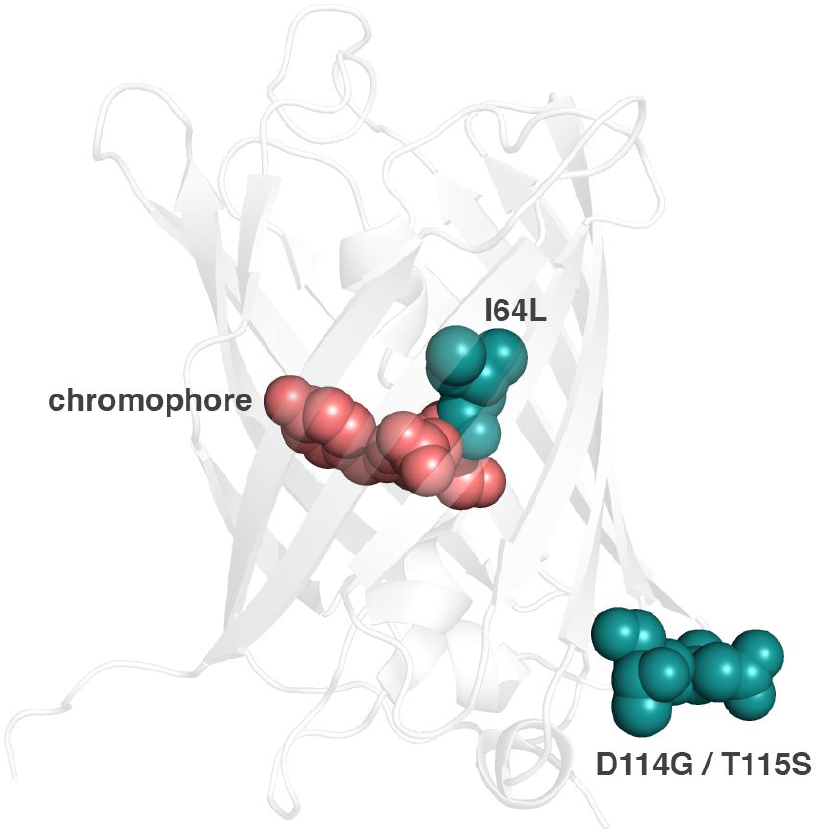
Location of the chromophore and mutations I64L, D114G and T115S in the structure of KillerRed.

## Conclusions

The KillerRed variant described in this work is a promising template for further directed evolution to create a new generation of genetically encodable photosensitizers with green fluorescence. Unlike other KillerRed spectral variants developed to date, the KillerRed I64L/D114G/T115S mutant forms a tyrosine-based chromophore and exhibits fluorescence in the green region of the spectrum.

## Acknowledgements

This work would not have been published without the generous support and the strict publication requirements of the Russian Foundation for Basic Research grant 18-04-01173. KSS is supported by the president fellowship 075-15-2019-411. Experiments were partially carried out using the equipment provided by the Institute of Bioorganic Chemistry of the Russian Academy of Sciences Сore Facility (CKP IBCH; supported by Russian Ministry of Education and Science Grant RFMEFI62117X0018).

